# Long-term vegetation change in protected calcareous fens is driven by land-use change and management abandonment

**DOI:** 10.64898/2026.07.20.739508

**Authors:** Miriam Diez, Alexander Peringer, Isabell Braunstein, Andreas Schweiger

**Affiliations:** University of Hohenheim, Institute of Landscape and Plant Ecology, Department of Plant Ecology, Stuttgart, Germany; University of Hohenheim, Institute of Agricultural Sciences in the Tropics, Department of Ecology of Tropical Agricultural Systems, Stuttgart, Germany; Nürtingen-Geislingen University, Landscape Ecology and Resource Conservation, Nürtingen, Germany

**Keywords:** Historical land use, land-use legacy, traditional management, vegetation resurvey, species turnover, beta diversity, peatlands, biodiversity conservation

## Abstract

**Aims:** Historical land use has shaped plant diversity in Central Europe for centuries, and many plant communities of high conservation value have been maintained by long-term traditional management. This study assesses how changes in land use over the past century have affected plant community composition and vegetation-based ecological indicator values in the management-dependent mire community *Primulo-Schoenetum ferruginei*.

**Location:** South-western Germany.

**Methods:** We conducted repeated vegetation surveys at multiple sites, comparing historical records with recent surveys spanning up to 97 years. Changes in species composition were analysed using multivariate approaches, and shifts in habitat conditions were assessed using indicator values for nutrients, temperature, light and mowing frequency. Information on historical management was compiled from original vegetation sources, while current management data were obtained from responsible conservation authorities and regional administration.

**Results:** Fen communities showed pronounced temporal changes in species composition. Community turnover was significantly associated with shrub encroachment, mowing frequency, hay transfer and early mowing. In addition, species-based indicator values suggested increasing nutrient availability and temperature affinity of the vegetation, alongside decreasing light availability. Disturbance indicators pointed towards less frequent but more severe disturbance regimes in contemporary compared to historical vegetation.

**Conclusions:** Our results demonstrate that management-dependent fen communities are highly sensitive to changes in land-use regimes, i.e. disturbance frequency and severity that is indicated to have significantly changed in the protected areas in this study. These findings highlight the importance of maintaining traditional land management practices when aiming to maintain biodiversity in calcareous fens.

## Introduction

Human influence affects most natural landscapes, shaping ecosystems and altering their species composition (Plumptre *et al*. 2021). Agricultural use has a centuries-long history in Central Europe and has profoundly influenced the development of specific plant communities (Ellis *et al*. 2021). Traditional, low-intensity land management has played a key role in shaping and maintaining species-rich communities that provided habitats for many taxa (Brown & Kothari 2011). These practices not only shaped ecosystems but also redefined niches and interactions within plant communities, particularly in open-land systems such as calcareous fens.

However, the decline of traditional land use practices increasingly threatens biodiversity, contributing to the loss of specialists and protected species (Plieninger & Bieling 2012). Across formerly traditionally managed sites, current management often differs substantially from the historical regimes due to either management intensification or abandonment (Middleton *et al*. 2006; van Diggelen *et al*. 2006). Both processes can alter environmental conditions and reduce the diversity of specialist species historically associated with these habitats (Fischer *et al*. 2012; Uchida & Ushimaru 2015; Colom *et al*. 2021). These processes can also promote biotic homogenization by favouring generalist species (Sutcliffe *et al*. 2015).

Fen communities are particularly sensitive to such changes due to their reliance on specific management regimes, such as mowing, to maintain open conditions and prevent shrub encroachment (Tanneberger *et al*. 2022). Traditionally, many Central European calcareous fens were managed as low-intensity litter meadows (“Streuwiesen”), characterised by annual late-season mowing and biomass removal, which helped maintain open, species-rich vegetation (Tanneberger *et al*. 2022; Ross *et al*. 2019). Despite their ecological importance, long-term vegetation resurveys to quantify effects of land use change on vegetation dynamics remain uncommon, as historical vegetation records rarely provide consistent plot-level management metadata (Kapfer *et al*. 2016; Jandt *et al*. 2022).

To address this gap, we used the calcareous fen community *Primulo-Schoenetum ferruginei* as a model system to investigate long-term vegetation change and its association with shifts in land use and management. In calcareous fens, being characterized by specialist species that response sensitively to management practices and environmental change, shifts in mowing frequency or timing, or the cessation of use may alter habitat conditions in ways that affect the persistence of characteristic species (Billeter *et al*. 2007; Plieninger *et al*. 2015).

Specifically, we ask (i) how species composition and biodiversity in *Primulo-Schoenetum ferruginei* have changed over the past century, (ii) whether habitat conditions and land-use management have shifted over time, and (iii) whether variation in land-use management (e.g. mowing and shrub encroachment) is associated with differences in species composition.

## Methods

### Historical sources and site selection

Using *Süddeutsche Pflanzengesellschaften* (Oberdorfer 1992) as the primary guide, we identified historical vegetation surveys of *Primulo-Schoenetum ferruginei* and selected four representative calcareous fen sites in south-western Germany (Figure 1**Fehler! Verweisquelle konnte nicht gefunden werden**.; Appendix S1, Table S1). Historical vegetation records were available from published surveys conducted at Wurzacher Ried (Ilschner 1959), Egelsee near Gornhofen (Bertsch 1928), Federsee (Kuhn 1954), Mindelsee (Lang 1973). The original sources, including dissertations, explanatory notes to vegetation maps (where available), and related publications, were compiled and analysed. Only sites for which historical surveys provided sufficient floristic detail and relocatable plot information were included, constraining the pool of usable resurveys. Species lists and abundance data were extracted from these sources, and all species names were systematically verified and updated to current nomenclature standards using the Plants of the World Online database (POWO 2025).

**Figure 1.**
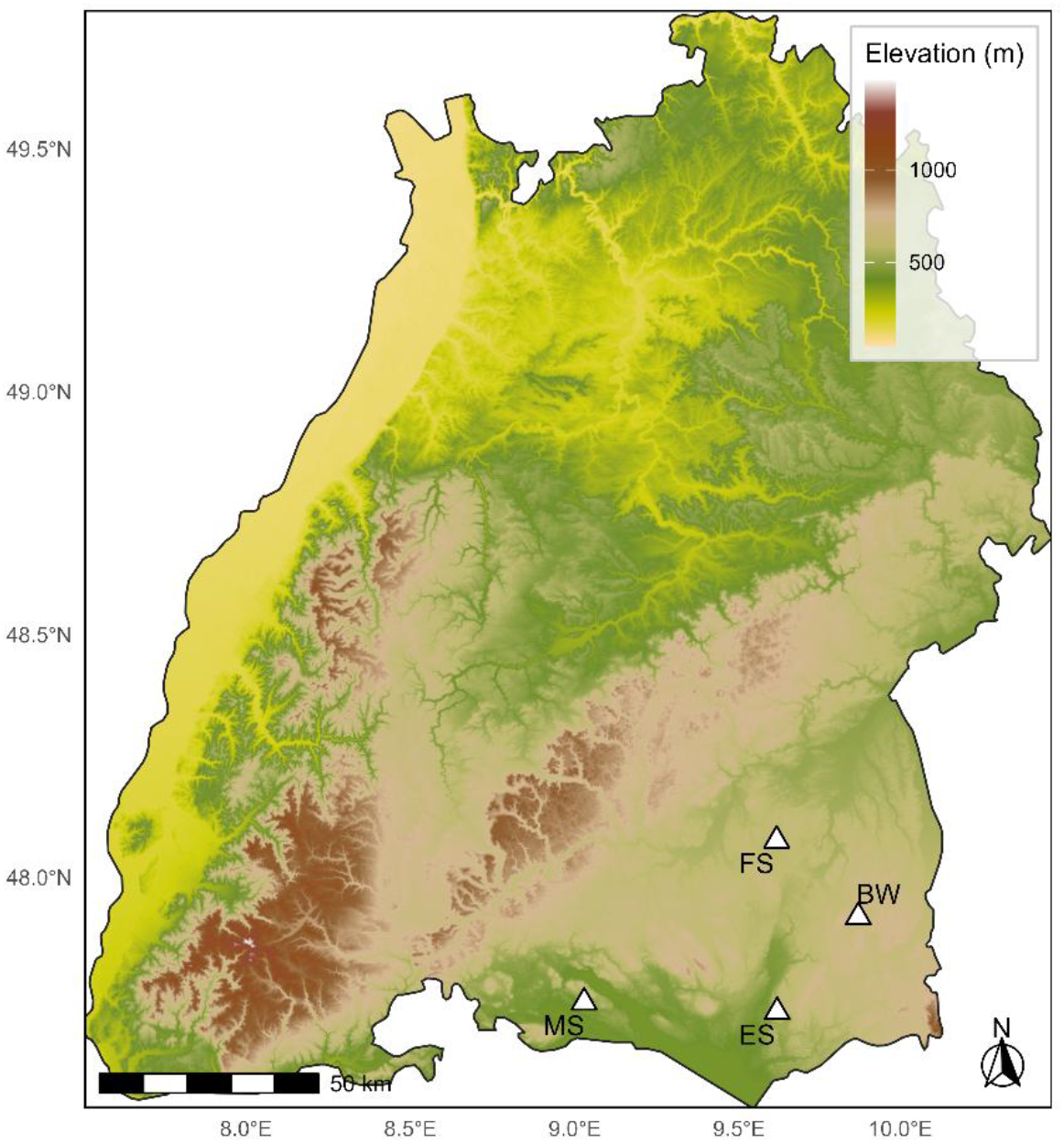
Elevation map of Baden-Württemberg (Germany) showing the four study sites of calcareous fens classified as *Primulo– Schoenetum ferruginei*. Elevation is derived from the SRTM digital terrain model. Site abbreviations: FS Federsee, BW Wurzacher Ried, MS Mindelsee, ES Egelsee.

In addition to floristic information, the historical sources and related regional literature were evaluated for information on former land use and management. Across all investigated calcareous fen sites, the literature consistently describes traditional low-intensity litter meadow management (“Streuwiesen”), typically involving annual late-season mowing and removal of cut biomass (Kuhn 1954; Ilschner 1959; Günzl 1989). As *Primulo-Schoenetum ferruginei* is closely linked to this management regime, all selected sites shared similar historical land-use conditions, while recent management continuity differed among sites. Information on recent management continuity and land-use change was compiled at plot level from conservation management plans, regional documentation, and communications with local conservation authorities and reserve managers.

### Recent vegetation surveys

Vegetation surveys were conducted at four calcareous fen sites (see Appendix S1 for site details). Survey plots measured 5 × 5 m. Within each plot, all vascular plant species were recorded and their percentage cover estimated. Although vegetation layers were recorded in the field where present, historical fen relevés generally did not distinguish between layers. To ensure comparability, all analyses were conducted at the plot level. Coordinates and elevation were documented for each plot. Species identification was verified using the *Rothmaler* flora (Jäger *et al*. 2017) where necessary, and nomenclature was standardised according to the Plants of the World Online database (POWO 2025). Recent plots were located within the historical study areas but could not always be matched to exact historical coordinates, resulting in a quasi-resurvey design. The number of historical and recent plots differed among sites because the availability of suitable historical vegetation records varied considerably (Appendix S1, Table S1).

### Analyses

To enable comparison between historical and contemporary vegetation surveys, plant cover values were standardised. Contemporary surveys recorded percentage cover as percentage values, whereas historical surveys used the Braun–Blanquet scale with five classes: 1 (1–5%), 2 (5–25%), 3 (25–50%), 4 (50– 75%), 5 (75–100%) (Braun-Blanquet 1932). To harmonise both datasets, Braun-Blanquet classes were converted to class midpoints (1 = 3%, 2 = 15%, 3 = 37.5%, 4 = 62.5%, 5 = 87.5%), enabling consistent comparisons while minimising information loss.

#### Alpha-diversity

We calculated the Shannon index (H′) per plot from species-by-plot abundance matrices. Differences between historical (“old”) and recent (“new”) surveys were tested using Wilcoxon rank-sum tests. P-values were adjusted using the Benjamini–Hochberg (BH) procedure, and medians as well as median differences are reported. Analogous analyses were performed on plot-level species richness; site-level summaries are provided in the Appendix S1, Table S3).

#### Beta-diversity

Presence–absence community matrices were compiled and Jaccard dissimilarities were calculated from binary data using vegan::vegdist. Overall differences in species composition between historical and recent surveys were assessed using PERMANOVA (adonis2, 999 permutations), with permutations restricted within sites (strata = Site) to account for spatial clustering. Homogeneity of multivariate dispersion was evaluated with PERMDISP (betadisper, bias.adjust = TRUE), based on 999 permutations. To disentangle components of β-diversity, total Sørensen dissimilarity was partitioned into turnover (β_SIM_) and nestedness (β_SNE_) using the betapart package. To characterise species contributing to temporal turnover, species occurrence frequencies (% of plots occupied) were compared between historical and recent surveys. For each species, frequency changes were calculated as the difference between survey periods. Significance of changes in occurrence frequency was assessed using Fisher’s exact tests, and p-values were adjusted using BH procedure. Species with positive frequency changes were classified as “winners”, species with negative frequency changes as “losers”. Dissimilarities within survey periods (old–old vs. new–new) were compared using unpaired Wilcoxon rank-sum tests, and p-values were Benjamini– Hochberg-adjusted across vegetation types and β-components.

#### Ordination and land-use correlates

Non-metric multidimensional scaling (NMDS) was performed using metaMDS (*k* = 2; Bray–Curtis dissimilarities; trymax = 100) on abundance data based on percentage cover and Braun–Blanquet midpoint values. Final stress values are reported for the ordinations. To assess the relationship between community composition and land-use variables, we conducted a distance-based redundancy analysis (dbRDA; capscale with Bray–Curtis dissimilarities). To assess whether variation in recent land-use management was associated with differences in species composition, plot-level land-use predictors (mowing frequency, timing of mowing, afforestation, shrub encroachment and removal of cut material) were z-standardised prior to analysis. Site was included as a conditional term (Condition (Site)) to account for among-site variation. We report global and marginal permutation tests (999 permutations with blocks defined by Site), variance inflation factors (VIFs), and forward selection (ordiR2step) constrained by the adjusted R^2^.

#### Red-List categories

Species were matched to the German Red List (BfN 2018), and threat categories were converted to a numeric scale ranging from 10 (extinct) to 3 (least concern); with non-assignable categories treated as NA. For each plot, we calculated both the mean Red-List category and the number of Red-List species (categories V-0). Differences between historical and recent surveys were tested using Wilcoxon rank-sum tests. P-values were adjusted using the Benjamini–Hochberg procedure. Descriptive summaries (means, medians, *SD, IQR*) and test results are provided in Appendix S1. Higher numeric values indicate higher threat status.

#### Ellenberg indicator values and disturbance metrics

Ellenberg indicator values (L, T, F, R, N) and species-based disturbance indicator values (disturbance severity, disturbance frequency, mowing frequency, grazing pressure and soil disturbance; (Tichý *et al*. 2023)) were matched to species occurrences and averaged at the plot level using unweighted means. These variables represent ecological indicator values associated with species occurrences and do not constitute direct measurements of site management. Temporal differences between historical and recent surveys were tested for each variable using Wilcoxon rank-sum tests with Benjamini–Hochberg (BH) correction. For visualisation, values were z-standardised within variables and ordered by the absolute median change between time periods.BH-corrected significance levels were indicated by asterisks (Figure 4; Appendix S1, Table S10).

#### Software

Analyses were conducted in R 4.3.2. Key packages included *vegan* 2.6-x, *betapart* 1.5-x, *permute* 0.9-x, *ggplot2* 3.5-x, and *dplyr* 1.x and *tidyr* 1.x. Fixed random seeds (e.g. set.seed(123) for NMDS and set.seed(2) for forward selection) were used to ensure reproducibility.

## Results

### Alpha Diversity

Shannon diversity remained largely stable between historical and recent fen surveys (Figure 2). Median Shannon diversity was slightly lower in recent plots (median old = 1.91, median new = 1.81; Δ = −0.11), although this difference was not significant (Wilcoxon test, *p*_*adj*_ = 0.341). Overall median plot-level species richness remained unchanged between historical and recent surveys (median = 18 species in both periods), although variability was higher in recent plots (IQR = 13.5) than in historical plots (IQR = 8). Species richness was generally stable across sites, with a significant decrease at Mindelsee (*p*_*adj*_ = 0.037) and a marginal decrease at Wurzacher Ried (*p*_*adj*_ = 0.057; Appendix S1, Figure S1).

**Figure 2.**
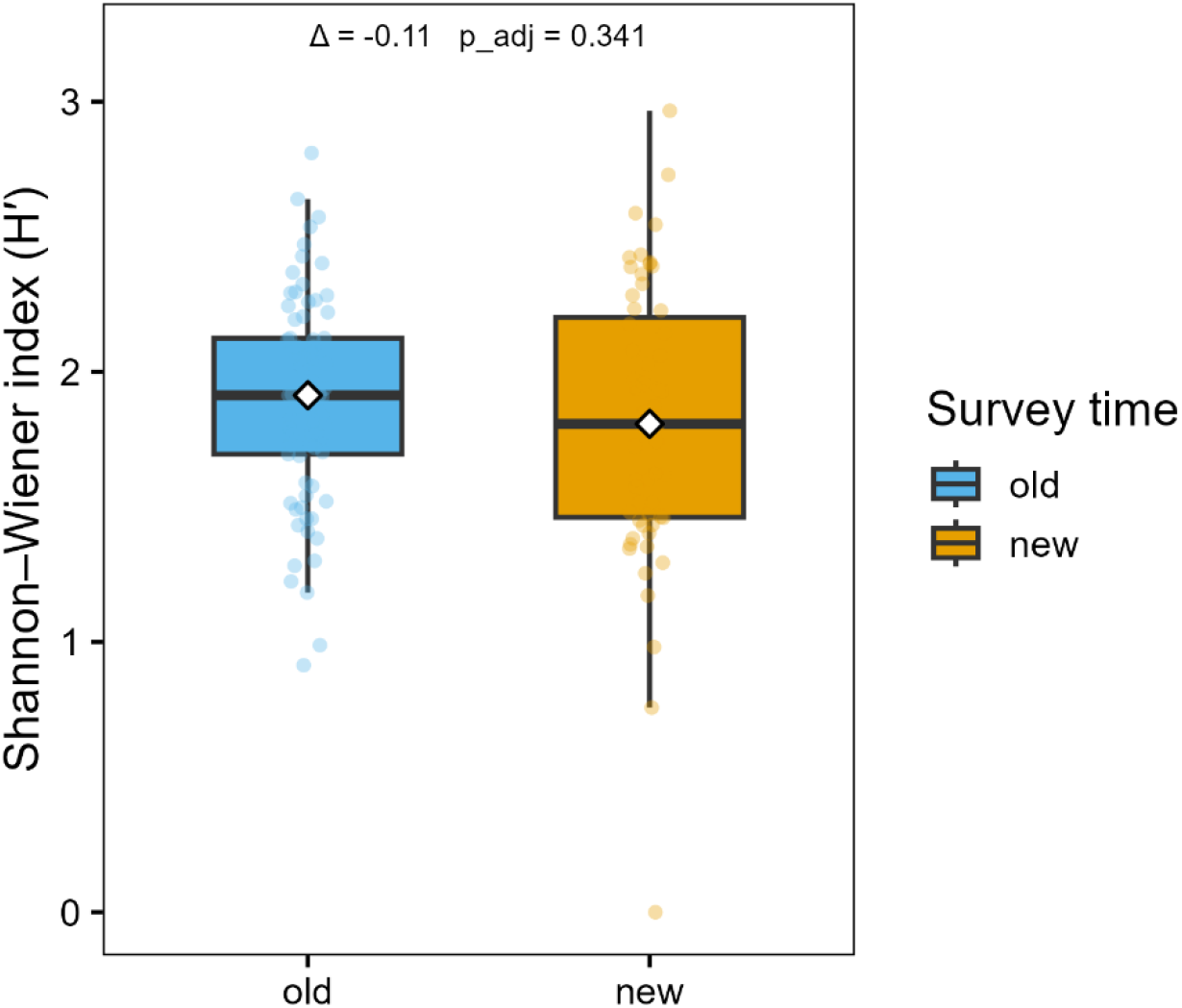
Shannon–Wiener diversity index (H′) in historical (n = 77) and recent (n = 59) surveys of calcareous fen plots. White diamonds indicate group medians. Δ and BH-adjusted p-values (*p*_adj_) are reported above the boxplots.

### Beta diversity: overall differences and turnover

Species composition differed significantly between historical and recent surveys. PERMANOVA based on Jaccard dissimilarities showed that survey period explained a small but significant proportion (2.3 %) of the variation in species composition (*F* = 3.103, *p* = 0.001). Community heterogeneity (distance to group centroids) did not differ significantly between historical and recent surveys (*F* = 2.29, *p* = 0.13; Appendix S1, Figure S2). Partitioning of Sørensen dissimilarities indicated that temporal *β*-diversity was primarily driven by species turnover rather than nestedness (Appendix S1, Figure S3, Table S5). Turnover was significantly lower among recent plots (*p*_adj_ < 0.001), while nestedness contributed little to overall dissimilarity and did not differ significantly between survey periods (*p*_adj_ = 0.052). Species-level frequency analyses further showed changes in the occurrence of individual species between historical and recent surveys (Appendix S1, Table S9). The largest declines were observed for *Tofieldia calyculata* (80.5% to 1.7% of plots), *Schoenus ferrugineus* (93.5% to 39.0%), *Parnassia palustris* (64.9% to 15.3%) and *Primula farinosa* (58.4% to 15.3%). In contrast, *Lythrum salicaria* increased from 2.6% to 42.4% of plots, *Carex flava* from 2.6% to 39.0%, and *Filipendula ulmaria* from 5.2% to 35.6%.

### Ordination and land-use correlates

NMDS based on Bray–Curtis dissimilarities (*k* = 2) summarized abundance-based compositional gradients (stress = 0.201; Appendix S1, Figure S4). Historical and recent surveys occupied partially distinct regions of ordination space, indicating directional changes in community composition over time. Distance-based RDA conditioned on Site indicated that land-use variables significantly explained variation in fen species composition (global model: *F* = 4.61, *p* = 0.001; adjusted *R²* = 0.10). Marginal effects revealed significant associations between species composition and mowing frequency, shrub encroachment, hay transfer, early mowing and afforestation (Table 1).

**Table 1.** Marginal effects of plot-level land-use predictors in the dbRDA model based on Bray–Curtis dissimilarities for calcareous fen plots, conditioned on Site. Reported are degrees of freedom (Df), sums of squares, F-statistics, permutation p-values based on 999 permutations, and Benjamini–Hochberg adjusted p-values (padj).

| Predictor | Df | Sum of squares | F | p | $p_{\text{adj}}$ |
| --- | --- | --- | --- | --- | --- |
| Early mowing | 1 | 1.213 | 4.119 | 0.001 | 0.001 |
| Mowing frequency | 1 | 1.544 | 5.242 | 0.001 | 0.001 |
| Afforestation | 1 | 0.892 | 3.029 | 0.005 | 0.005 |
| Shrub encroachment | 1 | 1.642 | 5.574 | 0.001 | 0.001 |
| Hay transfer | 1 | 1.409 | 4.785 | 0.001 | 0.001 |

### Red-List categories

Mean Red-List categories per plot were significantly higher in historical than in recent fen surveys (*p*_adj_ < 0.001), showing a higher representation of threatened species in the historical communities (Figure 3; Appendix S1, Table S8). Consistent with this pattern, the number of Red-List species per plot was significantly lower in recent surveys (Wilcoxon rank-sum test, p < 0.001). Across all surveys, the total number of recorded Red-List species declined from 82 in historical surveys to 57 in recent ones.

**Figure 3.**
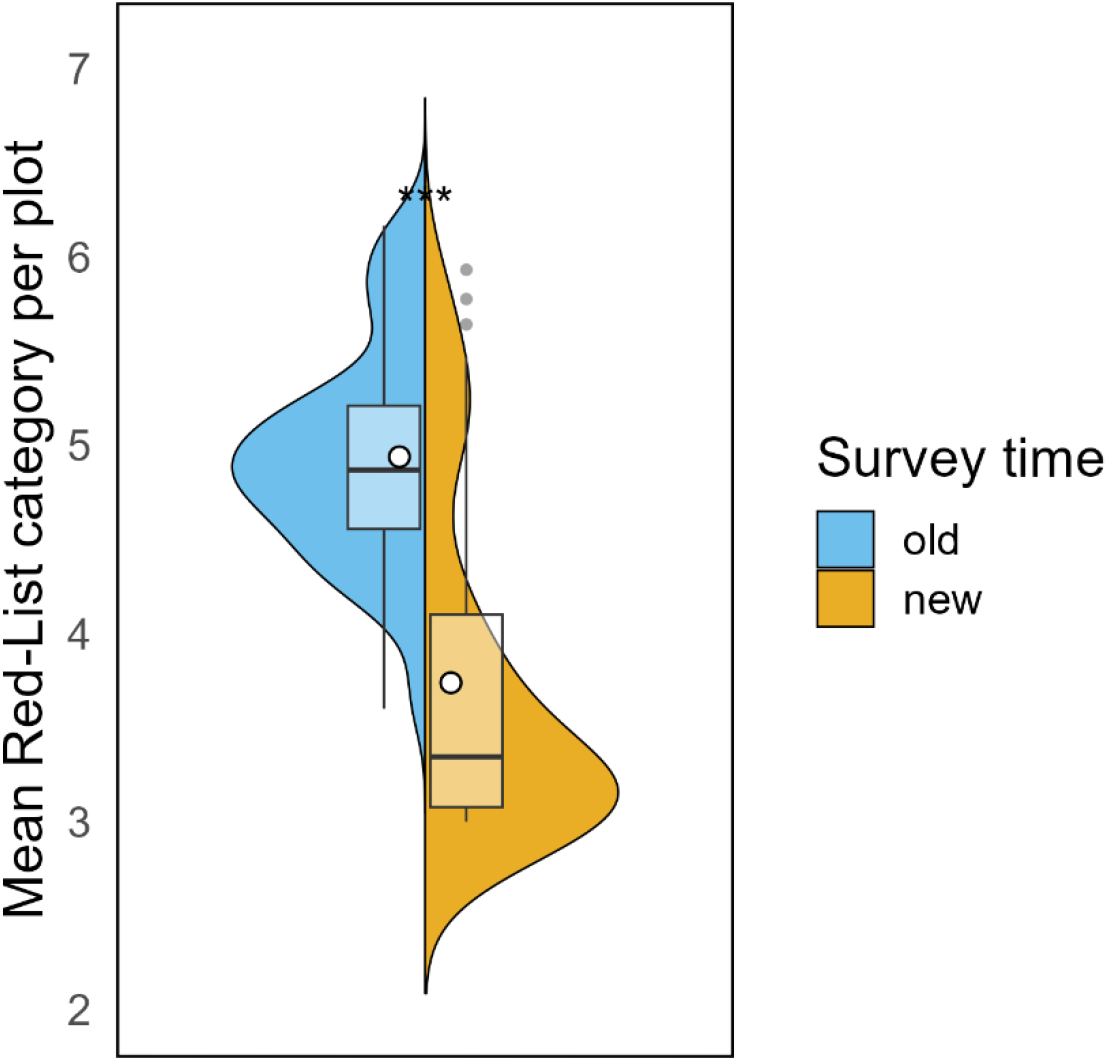
Split-violin plots showing mean Red List category per plot (higher values indicate higher threat status) in historical (n = 77) and recent (n = 59) surveys of calcareous fens. White points denote group means. Wilcoxon rank-sum tests revealed significantly lower mean threat status in recent plots (*p*_adj_ < 0.001).

### Ellenberg-type indicator values and disturbance metrics

Several Ellenberg-type indicator values and disturbance metrics shifted significantly between historical and recent surveys (Figure 4; Appendix S1, Table S10). Mean nutrient indicator values increased from 2.56 to 3.92 and mean temperature indicator values from 4.61 to 4.93, whereas mean light values decreased from 7.64 to 7.16 (all p_adj_ < 0.001). In contrast, moisture and reaction values did not differ significantly between survey periods. Species-based disturbance indicators are reported separately: disturbance severity increased from 0.38 to 0.46 and soil disturbance from 0.14 to 0.15, whereas disturbance frequency decreased from 1.08 to 0.92. Mowing frequency and grazing pressure did not differ significantly between survey periods.

**Figure 4.**
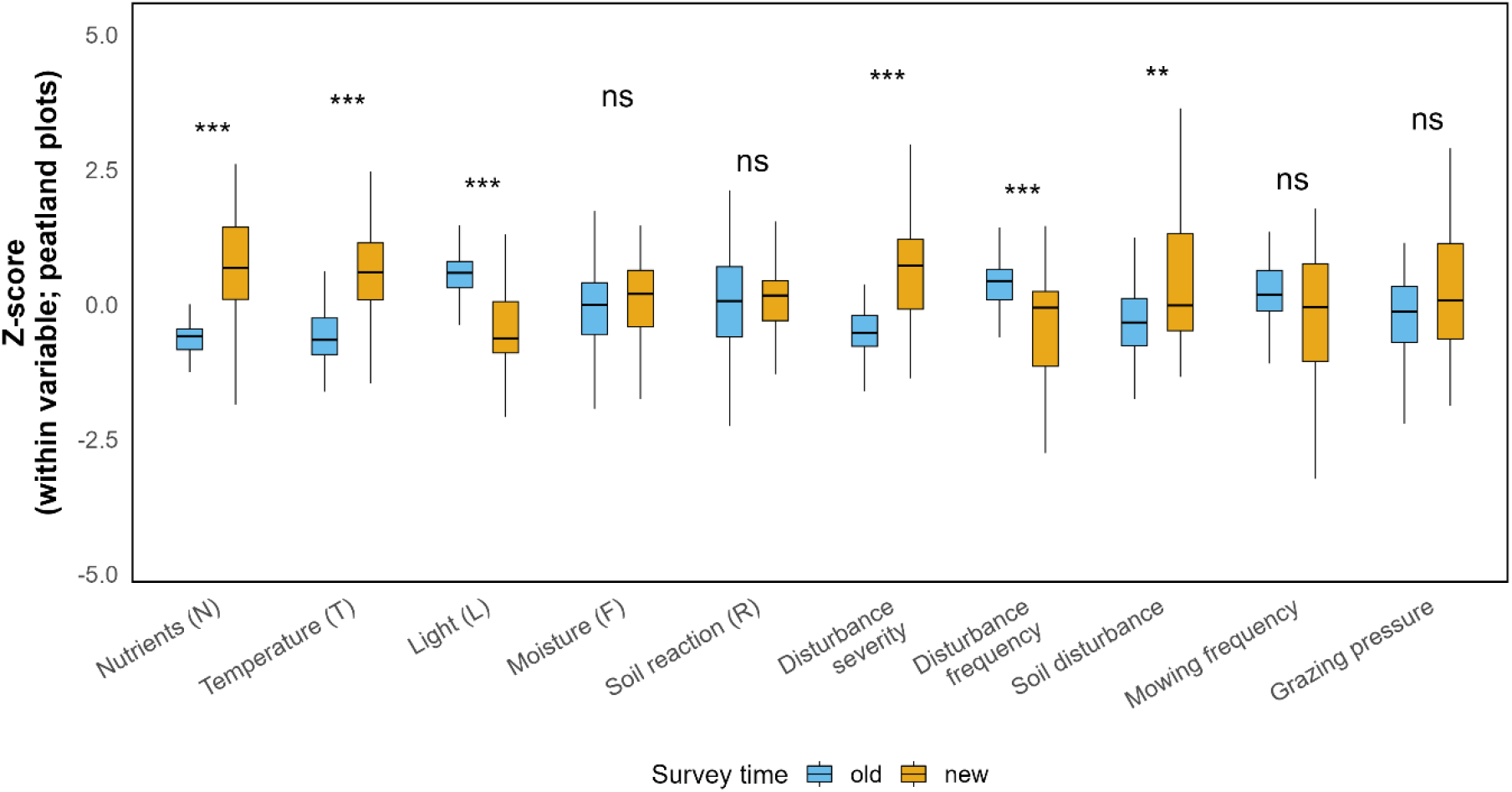
Temporal shifts in Ellenberg-type indicator values and disturbance metrics in historical (n = 77) and recent (n = 59) fen surveys. Values represent z-standardised plot-level means. Variables are grouped into Ellenberg indicators and disturbance metrics and ordered by absolute median change between survey periods. Asterisks denote BH-adjusted Wilcoxon rank-sum tests (* *p* < 0.05, ** *p* < 0.01, *** *p* < 0.001; *ns* = not significant). Herb-layer indices were excluded.

## Discussion

Our resurvey revealed pronounced temporal changes in calcareous fen community composition. These changes were primarily compositional and dominated by species turnover, while species richness and Shannon diversity remained largely stable over time.

Although PERMANOVA explained only a small proportion of the variance (*R²* ≈ 0.02–0.03), such values are common in multivariate community analyses and indicate directional shifts in species composition rather than differences in within-group dispersion (Tanneberger *et al*. 2022; Anderson 2006). This compositional change was mainly characterized by turnover rather than nestedness (Tanneberger *et al*. 2022; Baselga 2010). However, turnover was lower among recent plots, indicating increasing compositional similarity and potential homogenisation of present-day fen communities. This highlights that relatively stable diversity metrics can mask substantial compositional turnover and shifts in conservation value. The relatively small effect sizes likely reflect substantial spatial heterogeneity among sites, with some plots remaining largely stable while others exhibited pronounced shifts associated with differences in local management history, hydrological conditions, and shrub encroachment. Such patterns align with biodiversity declines following the abandonment of traditional land use documented across semi-natural grasslands and fen ecosystems (Eriksson *et al*. 2002; Middleton *et al*. 2006; van Diggelen *et al*. 2006).

Temporal shifts in fen community composition were closely associated with changes in management regimes. Distance-based redundancy analysis (dbRDA) identified shrub encroachment, mowing frequency, hay transfer, and early mowing as significant predictors of community turnover (Figure 1; Appendix S1, Table S6). These findings align with evidence that the abandonment of calcareous fens promotes shrub encroachment and hydrochemical change, disadvantaging calciphilous, weakly competitive species (Middleton *et al*. 2006; Lamers *et al*. 2015). A broader synthesis likewise shows that the biodiversity and trajectories of calcareous fen communities are strongly influenced by management, with abandonment acting as a consistent trigger for shrub encroachment and shifts in water regimes and chemistry (van Diggelen *et al*. 2006). However, the resumption of traditional low-intensity mowing, particularly when combined with biomass removal, can redirect these trajectories (Ross *et al*. 2019). The results support the importance of low-intensity management for maintaining species-rich calcareous fen vegetation and limiting further compositional homogenisation (Stammel *et al*. 2003). Consistent with these patterns, disturbance metrics indicated higher disturbance severity and soil disturbance, alongside lower disturbance frequency, whereas mowing frequency and grazing pressure did not differ significantly between survey periods (Appendix S1, Table S10).

Beyond direct management effects, compositional change in calcareous fens furthermore reflects shifts in abiotic habitat conditions. Ellenberg-type indicator values (Tanneberger *et al*. 2022; Tichý *et al*. 2023) indicated increasing nutrient availability and temperature, reduced light availability, and shifts in openness and soil chemistry, but no consistent decline in moisture conditions across plots. In addition, mean Red-List categories were higher in historical fen plots, followed by a strong shift towards lower mean threat status in recent plots, indicating a reduced representation of highly threatened species (Figure 3; Appendix S1, Tables S8-S9).

Species-level analyses revealed pronounced declines of several characteristic species of the historical *Primulo-Schoenetum ferruginei* stands that formed the basis of this resurvey, including *Tofieldia calyculata* (L.) Wahlenb., *Schoenus ferrugineus* L., *Primula farinosa* L., *Pinguicula vulgaris* L. and Parnassia palustris L. (Appendix S1, Table S9). These taxa are commonly associated with species-rich calcareous fens (Hájek *et al*. 2006; Wheeler & Proctor 2000). Their simultaneous decline indicates a weakening of the characteristic floristic composition historically associated with the target community. The decline of several diagnostic species occurred despite largely stable plot-level species richness, indicating that substantial changes in community composition and conservation value can occur even when overall diversity remains relatively constant. In contrast, the marked increase of species such as *Holcus lanatus* L., *Filipendula ulmaria* (L.) Maxim., *Carex acutiformis* Ehrh., *Lythrum salicaria* L. and *Salix cinerea* L. is consistent with a shift towards more productive and later-successional vegetation states. Such shifts are characteristic of abandoned or insufficiently managed calcareous fens, where litter accumulation, reduced disturbance frequency, and shrub encroachment favour competitive tall-growing species at the expense of light-demanding specialists adapted to nutrient-poor conditions (Middleton *et al*. 2006; van Diggelen *et al*. 2006; Lamers *et al*. 2015). This interpretation is supported by previous work showing that nutrient enrichment, lower water tables and the cessation of traditional mowing can promote grasses, forbs and shrub or woodland species while reducing fen specialists (Hájek *et al*. 2006; Middleton *et al*. 2006; van Diggelen *et al*. 2006; Lamers *et al*. 2015). The replacement of characteristic fen specialists by competitive taxa is therefore consistent with the increase in nutrient indicator values and the management-related predictors identified in the dbRDA analyses.

Finally, we critically evaluate key methodological choices, sources of uncertainty, and broader factors that may influence the interpretation of our findings. Braun-Blanquet cover classes were converted to midpoints (Braun-Blanquet 1928), and calculated Ellenberg-type as well as disturbance metrics were calculated as unweighted plot means. While these simplifications may reduce sensitivity to subtle changes, they are unlikely to affect the main conclusions of this study. Because management histories between historical and recent surveys are only available as snapshots, intermediate trajectories cannot be reconstructed, limiting inference on the timing and continuity of change. Taxonomic harmonisation and indicator assignment were controlled through systematic name standardisation, fuzzy matching with manual checks, and curated datasets (BfN 2018; Tichý *et al*. 2023). In addition, the paired resurvey design enabled direct temporal comparisons, and non-significant tests of multivariate dispersion confirmed that the observed compositional shifts represent directional change rather than altered within-group heterogeneity (Anderson 2006).

Broader drivers such as nitrogen deposition may also contribute to long-term vegetation change (Verheyen *et al*. 2012). Regional climate warming is well documented (Lenoir *et al*. 2008), and the observed increases in EIV-Temperature are consistent with thermophilisation patterns reported from other vegetation resurveys (Kiebacher *et al*. 2023). Hydrological change may contribute locally, however, the absence of a consistent decline in EIV-Moisture argues against widespread peatland desiccation across the investigated plots.

Limitations of the study include its regional focus on south-western Germany, the restriction to calcareous fen communities, a moderate sample size, and comparatively low multivariate R² values, indicating that compositional change did not occur uniformly across sites. The results should therefore be interpreted as evidence from one exemplar fen system rather than as broadly generalisable effects across ecosystems. Nevertheless, convergence across β-diversity components (Baselga 2010), Red-List status, Ellenberg-type values, disturbance metrics, and dbRDA supports the conclusion that the detected trends reflect genuine ecological change rather than methodological artefacts.

Taken together, our findings indicate pronounced long-term changes in calcareous fen vegetation despite largely stable species richness in Shannon diversity. The observed turnover was characterised by the decline of diagnostic fen species and increasing compositional similarity among contemporary communities, suggesting ongoing changes in conservation value that are not captured by traditional diversity metrics alone. Changes in management regimes, particularly shrub encroachment and alterations in mowing practices, were closely associated with these shifts, while species-based indicator values further pointed towards increasing nutrient availability and reduced light availability. Although the identified predictors likely represent correlated components of land-use change rather than independent causal drivers, the combined evidence suggests that changes in management intensity and type are main drivers of fen community reassembly and species turnover, even within protected areas. These findings highlight the importance of maintaining appropriate low-intensity management to conserve characteristic calcareous fen vegetation and its associated biodiversity.

## Appendices

Additional supporting information may be found online in the Supporting Information section: Appendix S1. Supplementary materials.

## Notes

### Competing Interest Statement

The authors have declared no competing interest.

