## Supplemental Data 1 for "Long-term vegetation change in protected calcareous fens is driven by land-use change and management abandonment"

Applied Vegetation Science

#### Methods

Table S 1 Site-level summary of the investigated calcareous fen sites, including survey years, number of historical and recent plots, median elevation, and bibliographic sources of the historical vegetation records.

| **Site (abbr.)** | **Site (full name)** | **First survey year** | **Repeated survey year** | **Historical plots** | **Recent plots** | **Median elevation (m a.s.l.)** | **Source of historical data** |
|---|---|---|---|---|---|---|---|
| BW | Wurzacher Ried | 1959 | 2024 | 60 | 24 | 831 | Ilschner, G. (1959) |
| ES | Egelsee bei Gornhofen | 1928 | 2025 | 2 | 10 | 676 | Bertsch, K. (1928) |
| FS | Federsee | 1954 | 2024 | 9 | 15 | 726 | Kuhn, L. (1954) |
| MS | Mindelsee | 1973 | 2025 | 6 | 10 | 529 | Lang, G. (1973) |

Table S 2 Overview of historical and recent management regimes across the investigated calcareous fen sites. Historical management was consistently characterised by traditional low-intensity litter meadow use ("Streuwiesen") with annual mowing and biomass removal, whereas recent management continuity differed among sites, ranging from continued conservation mowing to management abandonment.

| **Site** | **Historical management** | **Recent management** |
|---|---|---|
| Bw | annual litter mowing | mixed management continuity |
| ES | annual litter mowing | abandoned |
| FS | annual litter mowing | mixed management continuity |
| MS | annual litter mowing | unchanged |

### Alpha Diversity

Table S 3 Plot-level species richness in the investigated calcareous fen sites, reported as medians for historical (“old”) and recent (“new”) surveys. Differences between survey periods were tested using Wilcoxon rank-sum tests with Benjamini–Hochberg adjusted p-values (padj).

| Site | n_old | n_new | median_old | median_new | delta_median | p_adj |
|---|---|---|---|---|---|---|
| BW | 60 | 24 | 16 | 17 | 1 | 0.677 |
| ES | 2 | 10 | 17 | 7 | -10 | 0.361 |
| FS | 9 | 15 | 24 | 28 | 4 | 0.278 |
| **MS** | **6** | **10** | **22.5** | **17** | **-5.5** | **0.037** |

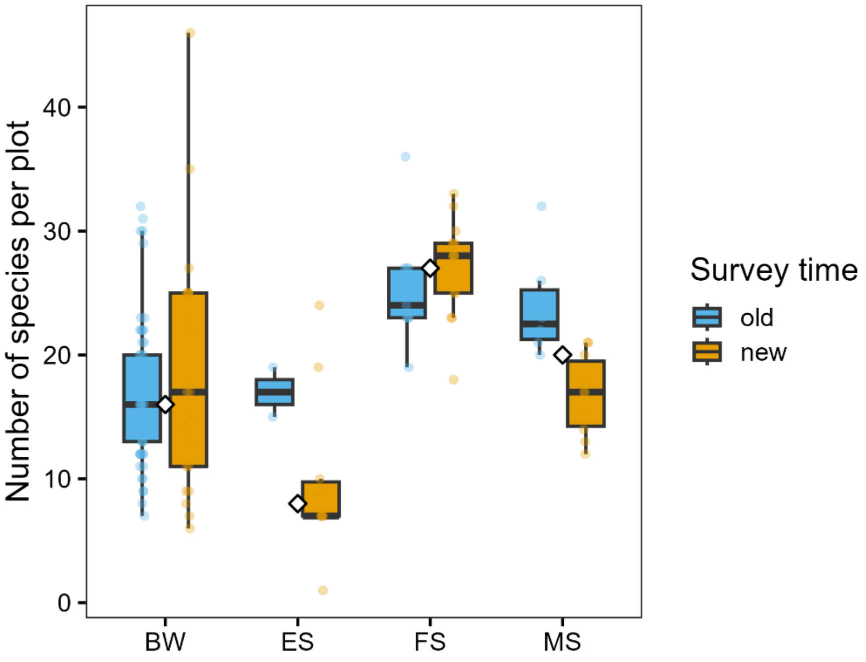


Figure S 1 Plot-level species richness in historical (“old”) and recent (“new”) calcareous fen surveys across the four study sites. Boxplots show medians, interquartile ranges, and whiskers (1.5 × IQR). White circles indicate group means, and grey points represent individual plots.

### Beta Diversity

Table S 4 Description of plot-level land-use and grouping variables used in the distance-based redundancy analyses (dbRDA), including variable type, coding, and interpretation. Information was compiled from conservation management plans, field observations, regional documentation, and communications with local conservation authorities and reserve managers. Analyses were performed for calcareous fen plots and conditioned on Site.

| **Variable** | **Type** | **Coding_or_unit** | **Description** |
|---|---|---|---|
| Mahd_früh | binary | 0/1 | Early mowing present this year |
| Mahd | integer | integer (events per year) | Number of mowings per year |
| Aufforstung | binary | 0/1 | Afforestation occurred |
| Verbuschung | binary | 0/1 | Shrub encroachment occurred |
| Mahdgut_ue | binary | 0/1 | Hay transfer performed |
| survey.time | factor | old/new | Historical vs. recent survey |
| Site | factor | levels = sites | Spatial block (permutation strata) |
| Plot | id | - | Permanent plot identifier |

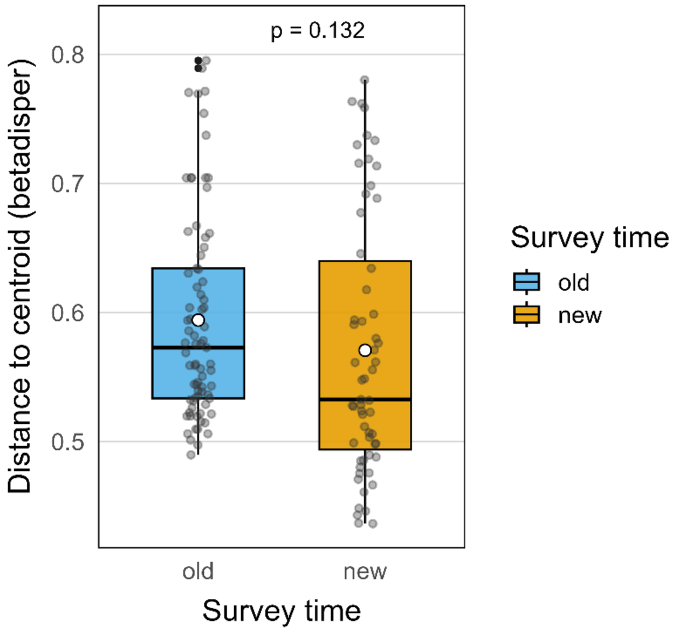


Figure S 2 Community heterogeneity (distance to group centroid, betadisper) of calcareous fen plots across survey times (“old” vs. “new”). Boxplots show medians, interquartile ranges, and whiskers (1.5 × IQR). White circles indicate group means, and grey points represent individual plot distances. Differences in multivariate dispersion between historical and recent surveys were not statistically significant (p = 0.132).

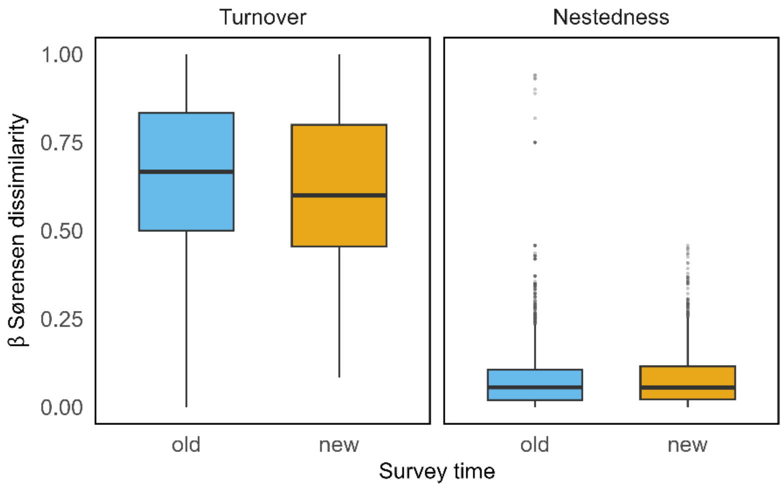


Figure S 3 Distribution of pairwise Sørensen dissimilarity components among calcareous fen plots within historical (“old–old”) and recent (“new–new”) surveys. Dissimilarity was partitioned into turnover (βSIM) and nestedness (βSNE) components. Boxplots show the distributions of pairwise dissimilarities; corresponding median values and Wilcoxon rank-sum tests are reported in Table S5.

Table S 5 Median (interquartile range, IQR) of pairwise Sørensen dissimilarities, partitioned into turnover (βSIM) and nestedness (βSNE) components, for historical (n = 77) and recent (n = 59) calcareous fen surveys. Differences between survey periods were tested using Wilcoxon rank-sum tests. Adjusted p-values (padj) are based on the Benjamini–Hochberg correction.

| Component | Median old [IQR] | Median new [IQR] | Wilcoxon p | $p_{adj}$ |
|---|---|---|---|---|
| Turnover (βSIM) | 0.67 [0.50–1.00] | 0.60 [0.48–0.95] | <0.001 | <0.001 |
| Nestedness (βSNE) | 0.06 [0.00–0.09] | 0.06 [0.00–0.09] | 0.052 | 0.052 |

Table S 6 Variance Inflation Factors (VIF) for land-use predictors retained in the dbRDA model. Lower VIF values indicate low collinearity among predictors; values below 5 are generally considered acceptable.

| Predictor | VIF |
|---|---|
| Mowing frequency | 2.226 |
| Shrub encroachment | 2.004 |
| Hay transfer | 1.396 |
| Afforestation | 1.175 |
| Early mowing (this year) | 1.165 |

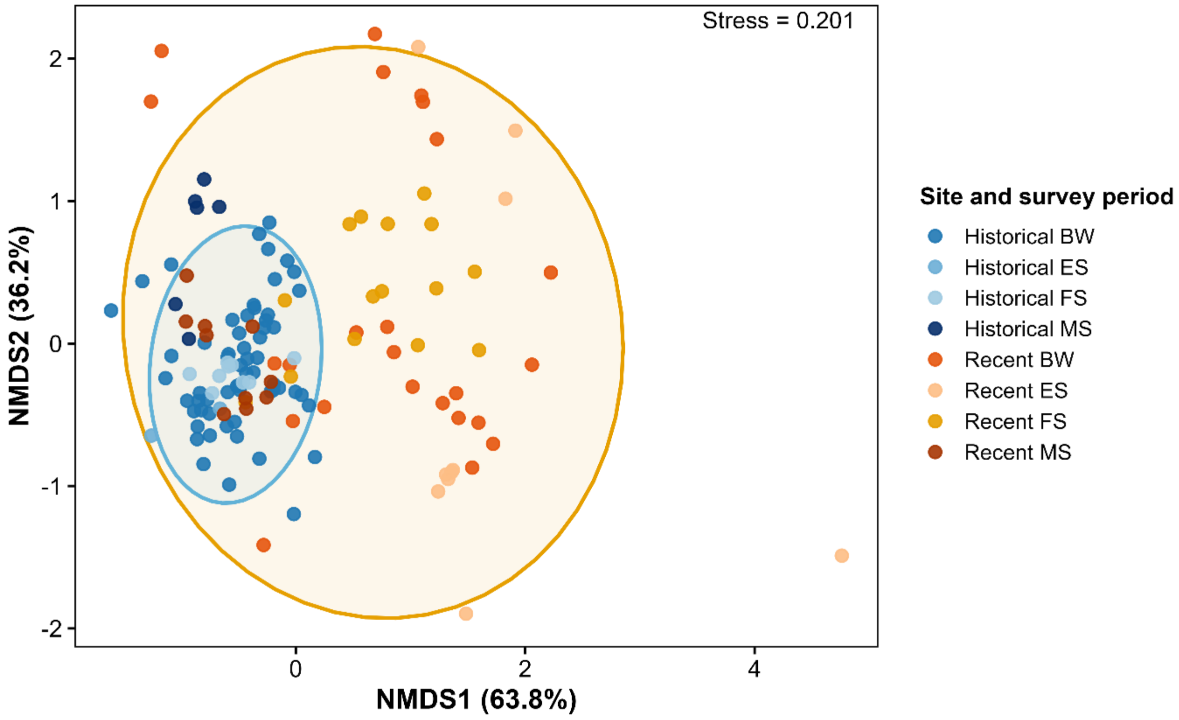


Figure S 4 Non-metric multidimensional scaling (NMDS) ordination of calcareous fen plots based on Bray–Curtis dissimilarities. Points represent individual plots coloured by site and survey period. Ellipses show 95% confidence regions for historical ("old") and recent ("new") surveys. Historical and recent communities occupied partially distinct regions of ordination space, indicating directional compositional change over time. Stress = 0.201.

### Red-List Category

Table S 7 Mapping of Red-List categories (BfN 2018) to numeric values used in the analyses. Higher numeric values indicate higher threat status. Data-deficient or non-assessed categories ("D", "nb", "kN", empty) were excluded (NA).

| **Original Red List code** | **Description (German)** | **Description (English)** | **Numeric value** |
|---|---|---|---|
| 0 | Ausgestorben oder verschollen | Extinct or missing | 10 |
| 1 | Vom Aussterben bedroht | Critically endangered | 9 |
| 2 | Stark gefährdet | Endangered | 8 |
| 3 | Gefährdet | Vulnerable | 7 |
| G | Gefährdung unbekannten Ausmaßes | Threatened, extent unknown | 6 |
| R | Extrem selten | Extremely rare | 5 |
| V | Vorwarnliste | Near threatened / warning list | 4 |
| * | Ungefährdet | Least concern | 3 |
| D, nb, kN, [empty] | Daten unzureichend / keine Bewertung | Data deficient / not assessed | NA |

Table S 8 Wilcoxon rank-sum tests comparing mean Red-List category per plot between historical ("old") and recent ("new") calcareous fen surveys. Reported are sample sizes, raw p-values (p), Benjamini–Hochberg adjusted p-values (padj), and significance codes (ns = not significant, *** = padj < 0.001).

| **n_old** | **n_new** | **p** | $p_{adj}$ | **Sig** |
|---|---|---|---|---|
| 77 | 59 | $3.41\times10^{-12}$ | $6.83\times10^{-12}$ | *** |

Table S 9 Species showing significant temporal changes in occurrence frequency between historical (n = 77) and recent (n = 59) calcareous fen surveys. Frequencies represent the percentage of plots in which a species occurred. Positive Δ Frequency values indicate increasing occurrence (winner species), whereas negative values indicate declining occurrence (loser species). Significance was assessed using Fisher's exact tests with Benjamini–Hochberg correction (padj < 0.05).

| **Species** | **Frequency old (%)** | **Frequency new (%)** | **Δ Frequency (%)** | **Direction** | **p** | **p adj** |
|---|---|---|---|---|---|---|
| *Tofieldia calyculata* (L.) Wahlenb. | 80,5 | 1,7 | -78,8 | Loser | <0.001 | <0.001 |
| *Schoenus ferrugineus* L. | 93,5 | 39 | -54,5 | Loser | <0.001 | <0.001 |
| *Parnassia palustris* L. | 64,9 | 15,3 | -49,7 | Loser | <0.001 | <0.001 |
| *Primula farinosa* L. | 58,4 | 15,3 | -43,2 | Loser | <0.001 | <0.001 |
| *Pinguicula vulgaris* L. | 49,4 | 8,5 | -40,9 | Loser | <0.001 | <0.001 |
| *Carex lepidocarpa* Tausch | 39 | 0 | -39 | Loser | <0.001 | <0.001 |
| *Leontodon hispidus* L. | 39 | 0 | -39 | Loser | <0.001 | <0.001 |
| *Salix repens* L. | 41,6 | 3,4 | -38,2 | Loser | <0.001 | <0.001 |
| *Succisa pratensis* Moench | 58,4 | 25,4 | -33 | Loser | <0.001 | 0.001 |
| *Linum catharticum* L. | 33,8 | 3,4 | -30,4 | Loser | <0.001 | <0.001 |
| *Trichophorum alpinum* (L.) Pers. | 29,9 | 0 | -29,9 | Loser | <0.001 | <0.001 |

| | | | | | | |
|---|---|---|---|---|---|---|
| *Molinia caerulea* (L.) Moench | 94,8 | 67,8 | -27 | Loser | <0.001 | <0.001 |
| *Drosera rotundifolia* L. | 29,9 | 3,4 | -26,5 | Loser | <0.001 | <0.001 |
| *Potentilla erecta* (L.) Raeusch. | 85,7 | 59,3 | -26,4 | Loser | <0.001 | 0.006 |
| *Frangula alnus* Mill. | 44,2 | 22 | -22,1 | Loser | 0.011 | 0.046 |
| *Luzula multiflora* (Ehrh.) Lej. | 22,1 | 0 | -22,1 | Loser | <0.001 | <0.001 |
| *Menyanthes trifoliata* L. | 28,6 | 8,5 | -20,1 | Loser | 0.004 | 0.025 |
| *Vaccinium oxycoccos* L. | 20,8 | 3,4 | -17,4 | Loser | 0.004 | 0.022 |
| *Centaurea jacea* L. | 15,6 | 0 | -15,6 | Loser | 0.001 | 0.008 |
| *Drosera intermedia* Hayne | 14,3 | 0 | -14,3 | Loser | 0.002 | 0.014 |
| *Polygala amara* L. | 14,3 | 0 | -14,3 | Loser | 0.002 | 0.014 |
| *Dactylorhiza incarnata* (L.) Soó | 15,6 | 1,7 | -13,9 | Loser | 0.007 | 0.033 |
| *Viola palustris* L. | 15,6 | 1,7 | -13,9 | Loser | 0.007 | 0.033 |
| *Rhynchospora alba* (L.) Vahl | 13 | 0 | -13 | Loser | 0.005 | 0.028 |
| *Carex lasiocarpa* Ehrh. | 10,4 | 0 | -10,4 | Loser | 0.010 | 0.045 |
| *Festuca ovina* L. | 10,4 | 0 | -10,4 | Loser | 0.010 | 0.045 |
| *Polygala amarella* Crantz | 10,4 | 0 | -10,4 | Loser | 0.010 | 0.045 |
| *Carex appropinquata* Schumach. | 0 | 10,2 | 10,2 | Winner | 0.006 | 0.029 |
| *Hypericum tetrapterum* Fr. | 0 | 10,2 | 10,2 | Winner | 0.006 | 0.029 |
| *Myosotis scorpioides* L. | 0 | 10,2 | 10,2 | Winner | 0.006 | 0.029 |
| *Viburnum opulus* L. | 0 | 10,2 | 10,2 | Winner | 0.006 | 0.029 |
| *Alnus glutinosa* (L.) Gaertn. | 0 | 11,9 | 11,9 | Winner | 0.002 | 0.014 |
| *Epilobium hirsutum* L. | 0 | 11,9 | 11,9 | Winner | 0.002 | 0.014 |
| *Festuca rubra* L. | 0 | 11,9 | 11,9 | Winner | 0.002 | 0.014 |
| *Prunus padus* L. | 0 | 11,9 | 11,9 | Winner | 0.002 | 0.014 |
| *Scutellaria galericulata* L. | 0 | 11,9 | 11,9 | Winner | 0.002 | 0.014 |
| *Vicia cracca* L. | 1,3 | 13,6 | 12,3 | Winner | 0.010 | 0.046 |
| *Sorbus aucuparia* L. | 0 | 13,6 | 13,6 | Winner | <0.001 | 0.007 |
| *Typha latifolia* L. | 0 | 13,6 | 13,6 | Winner | <0.001 | 0.007 |
| *Valeriana officinalis* L. | 2,6 | 18,6 | 16 | Winner | 0.002 | 0.014 |
| *Agrostis stolonifera* L. | 0 | 16,9 | 16,9 | Winner | <0.001 | 0.001 |
| *Lathyrus pratensis* L. | 0 | 18,6 | 18,6 | Winner | <0.001 | <0.001 |
| *Juncus subnodulosus* Schrank | 5,2 | 25,4 | 20,2 | Winner | <0.001 | 0.007 |
| *Phragmites australis* (Cav.) Trin. ex Steud. | 58,4 | 79,7 | 21,2 | Winner | 0.010 | 0.045 |
| *Deschampsia cespitosa* (L.) P.Beauv. | 5,2 | 27,1 | 21,9 | Winner | <0.001 | 0.004 |
| *Juncus effusus* L. | 0 | 23,7 | 23,7 | Winner | <0.001 | <0.001 |
| *Salix cinerea* L. | 0 | 25,4 | 25,4 | Winner | <0.001 | <0.001 |
| *Lotus pedunculatus* Cav. | 2,6 | 28,8 | 26,2 | Winner | <0.001 | <0.001 |
| *Carex acuta* L. | 0 | 28,8 | 28,8 | Winner | <0.001 | <0.001 |
| *Juncus articulatus* L. | 7,8 | 37,3 | 29,5 | Winner | <0.001 | <0.001 |

| | | | | | | |
|---|---|---|---|---|---|---|
| *Filipendula ulmaria* (L.) Maxim. | 5,2 | 35,6 | 30,4 | Winner | <0.001 | <0.001 |
| *Carex acutiformis* Ehrh. | 0 | 30,5 | 30,5 | Winner | <0.001 | <0.001 |
| *Galium uliginosum* L. | 11,7 | 42,4 | 30,7 | Winner | <0.001 | <0.001 |
| *Holcus lanatus* L. | 0 | 32,2 | 32,2 | Winner | <0.001 | <0.001 |
| *Lysimachia vulgaris* L. | 1,3 | 33,9 | 32,6 | Winner | <0.001 | <0.001 |
| *Galium palustre* L. | 3,9 | 37,3 | 33,4 | Winner | <0.001 | <0.001 |
| *Carex flava* L. | 2,6 | 39 | 36,4 | Winner | <0.001 | <0.001 |
| *Lythrum salicaria* L. | 2,6 | 42,4 | 39,8 | Winner | <0.001 | <0.001 |

### Ellenberg-Type Indicator Values

Table S 10 Temporal changes in Ellenberg indicator values and disturbance metrics for calcareous fen plots. Indicator values were aggregated as unweighted plot-level means. Differences between historical ("old") and recent ("new") surveys were tested using Wilcoxon rank-sum tests. Reported are medians, means, raw p-values (p), and Benjamini–Hochberg adjusted p-values (padj), including significance codes (ns, *, **, ***).

| **Variable** | **Median old** | **Median new** | **Mean old** | **Mean new** | **p** | $p_{adj}$ | **Sig** |
|---|---|---|---|---|---|---|---|
| Light | 7.654 | 7.196 | 7.637 | 7.164 | 0.000 | 0.000 | *** |
| Temperature | 4.594 | 4.895 | 4.612 | 4.932 | 0.000 | 0.000 | *** |
| Moisture | 8.096 | 8.193 | 8.049 | 8.160 | 0.122 | 0.153 | ns |
| Reaction | 5.908 | 5.981 | 5.855 | 5.867 | 0.680 | 0.680 | ns |
| Nutrients | 2.600 | 3.786 | 2.559 | 3.915 | 0.000 | 0.000 | *** |
| Disturbance Severity | 0.381 | 0.464 | 0.382 | 0.460 | 0.000 | 0.000 | *** |
| Disturbance Frequency | 1.098 | 1.001 | 1.084 | 0.917 | 0.000 | 0.000 | *** |
| Mowing Frequency | 0.467 | 0.444 | 0.470 | 0.420 | 0.152 | 0.169 | ns |
| Grazing Pressure | 0.185 | 0.189 | 0.184 | 0.192 | 0.056 | 0.079 | ns |
| Soil Disturbance | 0.135 | 0.141 | 0.135 | 0.149 | 0.001 | 0.001 | ** |
